# Transcription-coupled repair in *Drosophila melanogaster* is independent of the mismatch repair pathway

**DOI:** 10.1101/2020.04.07.029033

**Authors:** Lauri Törmä, Claire Burny, Viola Nolte, Kirsten-André Senti, Christian Schlötterer

**Affiliations:** Institut für Populationsgenetik, Vetmeduni Vienna, Vienna, Austria; Vienna Graduate School of Population Genetics, Vetmeduni Vienna, Vienna, Austria

**Keywords:** Transcription-coupled repair, *Drosophila melanogaster*, mutational strand bias, mismatch-repair, *spellchecker1*

## Abstract

Transcription-coupled repair (TCR) removes base damage on the transcribed strand of a gene to ensure a quick resumption of transcription. Based on the absence of key enzymes for TCR and empirical evidence, TCR was thought to be missing in *Drosophila melanogaster*. The recent demonstration of TCR in S2 cells raises the question about the involved genes. Since the mismatch repair (MMR) pathway serves a central role in TCR, at least in *Escherichia coli*, we studied the mutational signatures in flies with a deletion of the MMR gene *spellchecker1* (*spel1)*, a MutS homolog. Whole-genome sequencing of mutation accumulation (MA) lines obtained 7,345 new single nucleotide variants (SNVs) and 5,672 short indel mutations, the largest data set from an MA study in *D. melanogaster*. Based on the observed mutational strand-asymmetries, we conclude that TCR is still active without *spel1*. The operation of TCR is further confirmed by a negative association between mutation rate and gene expression. Surprisingly, the TCR signatures are detected for introns, but not for exons. We propose that an additional exon-specific repair pathway is masking the signature of TCR. This study presents the first step towards understanding the molecular basis of TCR in *Drosophila melanogaster*.

## Background

DNA continuously undergoes a large number of spontaneous chemical modifications leading to DNA damage (1, 2). The damaged bases can cause mutations, block DNA replication, and interfere with transcription (3). To repair some of these adducts, nucleotide excision repair (NER) removes the damaged strand at short distances from both sides of the lesion and a new strand is synthesized to fill the gap (4). NER has two pathways to recognize these lesions: the global genomic repair (GGR) and the transcription-coupled repair (TCR) (5). GGR scans the whole genome and the DNA damage is recognized by helix-distorting lesions (4). TCR detects the base damage from RNA polymerase stalling in actively transcribed DNA (4, 5) and leads to a mutational asymmetry between the strands (6). Transcription inhibition can be dangerous to a cell or even an organism (7, 8) so a quick resumption of transcription is vital. TCR is found in most bacterial species and many eukaryotes (9) and a defective pathway causes strong disease phenotypes, such as xeroderma pigmentosum and Cockayne’s syndrome in humans (10).

*Drosophila melanogaster* presents an interesting case where the GGR pathway for NER is present but TCR was thought to be missing (11, 12). The lack of fly homologs for genes required for TCR in other organisms — CSA/ERCC8 and CSB/ERCC6 — suggested that the pathway was lost during evolution (11). Furthermore, biochemical studies failed to detect TCR after UV-induced damage in *D. melanogaster* cell cultures (13, 14). In addition to the indirect evidence for TCR which is based on positive correlation of the compositional skews in introns (15) with expression (16), a recent study showed that TCR is operating in *Drosophila* S2 cells (17). This result raises the important question of how *Drosophila* is able to perform TCR when the key genes CSA and CSB are absent.

In *E. coli*, mismatch repair genes MutS and MutL are required for TCR (18). In yeast, genes required for NER interact with MMR genes (19) but MMR deficient cells are still performing TCR (20). In humans, MMR repair also interacts with NER (21) and evidence suggests that the pathway is involved in TCR of UV and oxidative damage (22). However, the issue remains controversial (23). Given the uncertainty about the functional basis of TCR in *Drosophila*, we determined the influence of the MutS homolog *spellchecker1* (*spel1*) on TCR in flies using MMR deficient mutation accumulation (MA) lines. The lack of MMR in MA lines is expected to result in a high number of mispaired bases. Such bases do not only lead to mutations but also cause local structural and dynamic distortions in the DNA structure (24) and are hotspots for DNA damage due to the higher susceptibility of unpaired bases to chemical modifications (25). For example, the loss of *msh2* in mice, *Trypanosoma brucei*, and *T. cruzi* increases oxidative damage of guanine by reactive oxygen species (26–28), which is repaired by TCR in murine cells (29).

Using a mismatch repair-deficient background, we find mutational asymmetries that are negatively associated with germline expression intensities, demonstrating functional TCR without *spel1*. Based on the absence of the TCR signatures in exons, we propose that an additional, exon-specific repair mechanism is operating.

## Results

We generated a *spel1* null mutant using CRISPR with guide RNAs targeting the 5’ and 3’ ends of the gene. Our mutant contained a double insertion of the template plasmid with the backbone of the vector (Supplementary Figure 1), a frequent event arising from the recombination of two plasmids into the locus (30). We propagated seven independent lines for 10 generations by brother-sister mating and identified 7,345 new single nucleotide variants (SNV) and 5,672 indels in females from these mutation accumulation lines. With 73.5% of the non-synonymous substitutions on the autosomes and 77.0% on the X chromosome, our data did not significantly deviate from the 75% expected under neutrality (31) (Fisher’s exact test (FET), p=0.4143 for the autosomes; FET, p=0.6573 for X).

The presence of TCR can be detected by mutational asymmetries between the transcribed and non-transcribed strands (6). The identification of mutational asymmetries is critically dependent on the correct null hypothesis. *Drosophila* introns have a skewed base composition, which depends on transcription levels (16). We confirmed that the fraction of thymines and cytosines on the transcribed strand is significantly negatively associated with expression in both ovaries (Wald test, OR=0.9977, p<2.2e-16 for thymines; Wald test, OR=0.9997, p=1.16e-13 for cytosines) and testes (Wald test, OR=0.9982, p<2.2e-16 for thymines; Wald test, OR=0.9994, p<2.2e-16 for cytosines) (Figure 1. a,b). We accounted for this by including the bias into the formulation of a null hypothesis for the expected number of mutations on the transcribed and on the non-transcribed strands. We calculated the expected bias with two different approaches: from the mutated genes and from a sample of genes with a similar expression as the mutated genes (see Methods). Both approaches produced highly consistent results.

**Figure 1.**
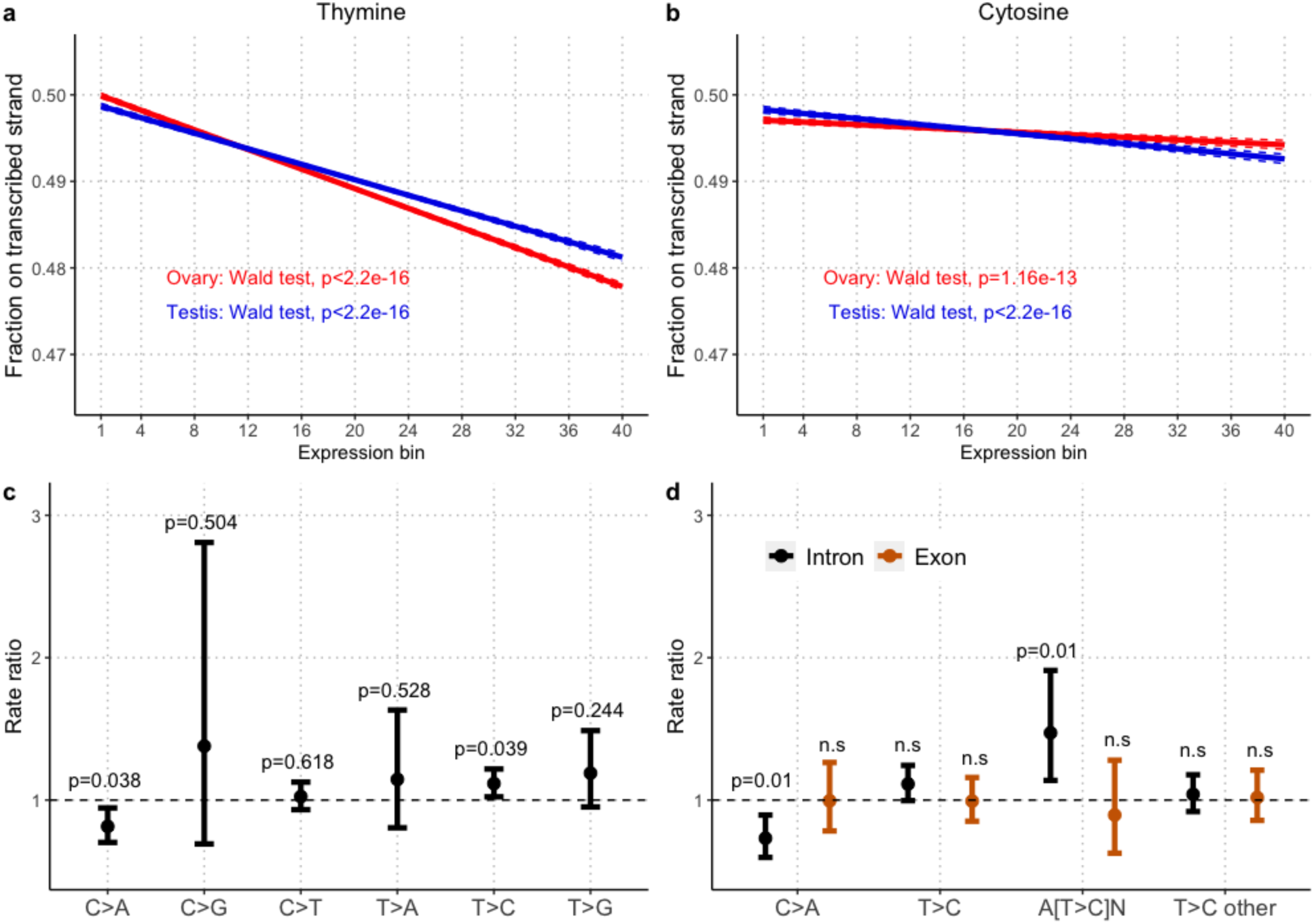
The interplay between transcription-associated base skews and mutational bias within genes. *Top*. Correlation between binned gene expression of 16,781 genes in ovaries (red) and testes (blue) and the fraction of a) thymines and b) cytosines on the transcribed strand. The regression line and its confidence interval are in plain and dotted lines respectively using the Generalized Linear Modeling framework. The p-values (Wald tests) correspond to the binned gene expression covariate. The intercept line of 1 indicates the absence of differences in rates of mutations on the transcribed and non-transcribed strands. The significance threshold is set to 5%. *Bottom*. Estimated rate ratios (RR) for different substitution types on the transcribed and non-transcribed strands in c) genes and partitioned for d) introns (black) and exons (orange). Adjusted p-values using the Benjamini-Hochberg procedure are reported and 95% Poisson confidence intervals are indicated by whiskers.

5,071 SNVs located in genes were used to test the ratio of bias-adjusted mutation rates on the transcribed and non-transcribed strands for every mutation type. Without TCR, a rate ratio (RR) of 1 is expected and the statistical significance can be determined with a Poisson test. After multiple testing correction, C>A mutations occurred less often (Poisson test, RR=0.82; 95% CI: 0.70-0.95, adjusted p-value=0.038), (Figure 1. c; Supplementary Table 1) and T>C mutations more often on the transcribed strand (Poisson test, RR=1.12, 95% CI: 1.02-1.22, adjusted p-value=0.039) (Figure 1. c; Supplementary Table 1). Similar results were obtained using the gene expression sampling scheme (see Methods, Supplementary Figure 2). Assuming that TCR is causing this bias, this implies that cytosine and adenine are more likely to experience base damage than other bases in MMR deficient flies.

DNA repair and damage processes can differ between exons and introns (32, 33). We therefore analyzed exons and introns separately. After excluding SNVs which overlapped both exon and intron annotations, C>A mutations occurred less often on the transcribed strand (RR=0.732, 95% CI: 0.597-0.895, adjusted p-value=0.014), but exonic C>A mutations did not (RR=0.995, 95% CI: 0.783-1.263, adjusted p-value=1) (Figure 1. d; Supplementary Table 2). Despite intronic T>C mutations occurring slightly more often on the transcribed strand, this was not significant (Poisson test, RR=1.113, 95% CI: 0.996-1.243, adjusted p-value=0.155). However, looking for the effect of the 5’ and 3’ bases flanking the mutation, we observed that the A[T>C]N context is exhibiting a significant strand bias in introns with a rate ratio of 1.472 (Poisson test, 95% CI: 1.138-1.910, adjusted p-value=0.01) but not in exons (Poisson test, RR=0.894, 95% CI: 0.627-1.280, adjusted p-value=0.866). No other contexts exhibited strand bias (Figure 1. d; Supplementary Table 2). Since the null hypothesis was not adjusted for triplet composition, we updated our null hypothesis to take into account the 5’ and 3’ flanking bases by performing a permutation test (see Methods) and obtained similar results. Intronic A[T>C]N mutations still exhibited a significant strand bias (permutation test, p=0.001) while exonic mutations did not (permutation test, p=0.215) (Supplementary Figure 3).

To confirm that the strand bias is caused by TCR, we tested for expression differences in genes containing C>A or A[T>C]N mutations. In the case of an active TCR, a correlation between strand asymmetry and gene expression is expected, because DNA damage on the transcribed strand is more likely to be detected in highly expressed genes. Thus, mutations arising from DNA damage on the transcribed strand should be found in lowly expressed genes. We used the FlyAtlas2 (34) expression data set from ovaries and testes as a proxy for the expression environment where the mutations occurred. Consistent with these predictions, we found that the genes with intronic C>A mutations on the transcribed strand have on average lower expression in both ovaries (one-sided Wilcoxon rank-sum test, adjusted p-value=0.022) and testes (one-sided Wilcoxon rank-sum test, adjusted p-value=0.022) than genes with intronic C>A mutations on the non-transcribed strand (Figure 2. a). As expected from the lack of strand bias, the expression level of genes with exonic C>A mutations were not different (Figure 2. a). Genes with context-dependent A[T>C]N mutations were not differentially expressed (Figure 2. b). This could be due to either a lack of power or because the expression data used does not reflect the expression environment where the base damage occurred.

**Figure 2.**
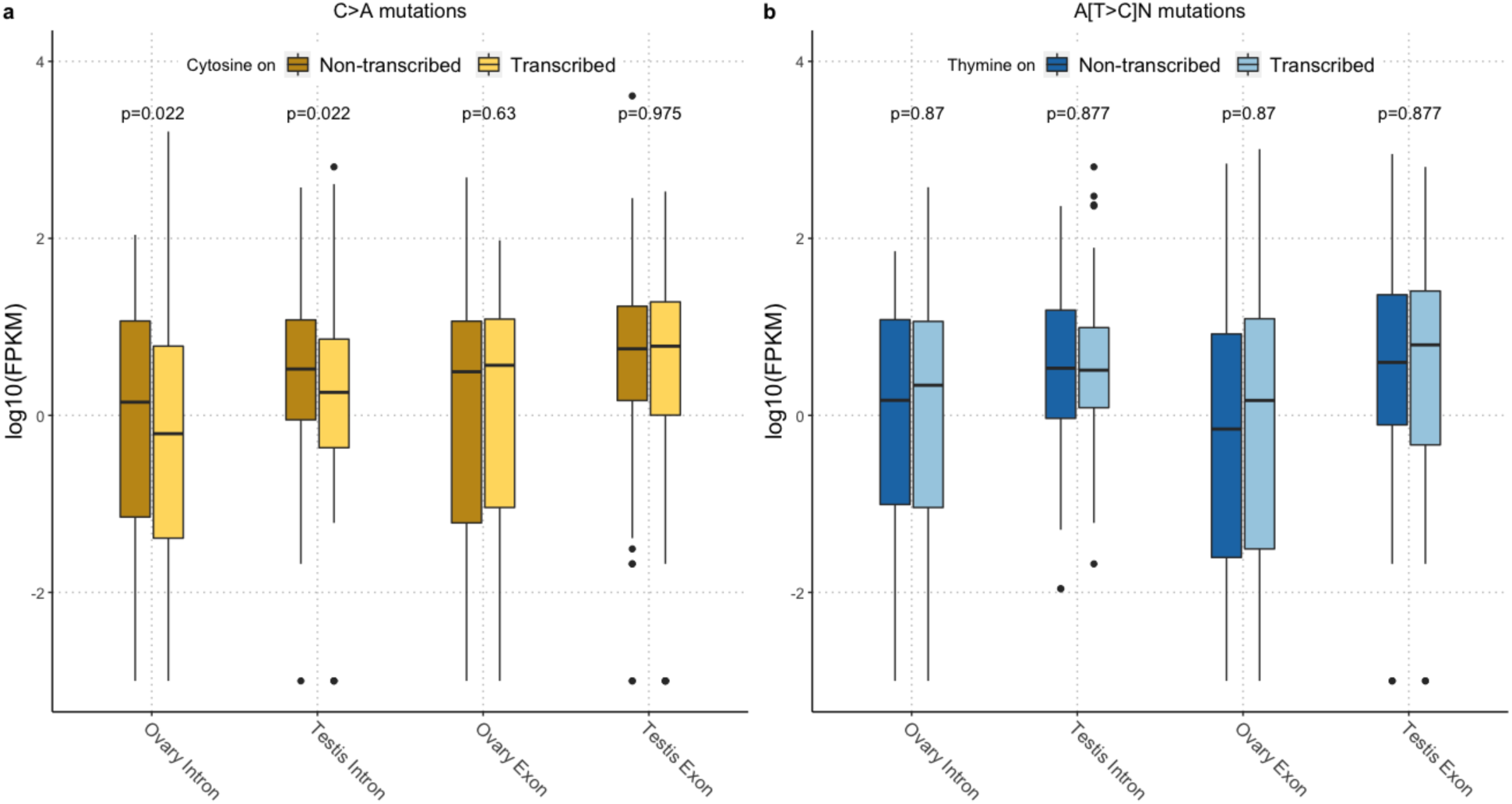
Introns with C>A mutations on the transcribed strand have lower expression levels. Boxplots of gene expression levels (log-10 transformed FPKM + *c*, with *c* = 0.001 to include genes not being expressed) of expressed mutated genes in ovaries and testes with a) C>A mutations and b) A[T>C]N mutations on the transcribed (light) and non-transcribed (dark) strands in exons and introns. The p-values are from one-sided (introns) or two-sided (exons) Wilcoxon rank-sum tests done on all genes and were adjusted per mutation type with the Benjamini-Hochberg procedure.

While the expression analysis suggests that TCR is responsible for the strand bias for C>A mutations, it is important to rule out the alternative explanation of a mutagenic effect of transcription on the non-transcribed strand. We used a randomization procedure (see Methods) to test if C>A mutations occur more frequently on the non-transcribed strand of highly expressed genes. Consistent with previous observations (16, 31), we found no evidence that transcription is mutagenic neither in testes (randomization test, p=0.611) nor in ovaries (randomization test, p=0.403) (Supplementary Figure 4) ruling it out as the source of the strand bias.

Based on the combined evidence, we conclude that TCR is operating in *Drosophila* biasing the C>A mutations and *spel1* is not required. Nevertheless, it is not clear why TCR signatures are only detected for introns, but not for exons. Two different explanations can account for the lack of mutational strand bias in exons for the C>A mutations: i) TCR requires *spel1* in exons or ii) an additional DNA repair mechanism is operating on exons, which erases the signal of TCR. The two explanations can be distinguished based on their different predictions for the relative mutation rates. MMR dependence for exons predicts an increased mutation rate for exons on the transcribed strand while the latter predicts a reduced exonic mutation rate for the non-transcribed strand. To test these hypotheses, we performed a permutation test while controlling for the triplet context in exons and introns to test for relative mutation rate differences. We found no evidence of elevated exonic mutation rate on the transcribed strand (permutation test, p=0.4762) (Supplementary Figure 5) showing that the lack of *spel1* does not cause the missing strand bias. We found signs - although nonsignificant - of reduced exonic mutation rate on the non-transcribed strand for C>A mutations (Supplementary Figure 5) (permutation test, p=0.067) suggesting that the lack of exonic strand bias may be caused by a favorable repair. The A[T>C]N did not show differences in the relative mutation rates on the transcribed (permutation test, p=0.475) or the non-transcribed strand (permutation test, p=0.126) (Supplementary Figure 6).

## Discussion

We demonstrated that TCR is independent of MMR in flies by uncovering TCR-induced mutational asymmetries in intronic C>A mutations in MMR deficient *D. melanogaster* mutation accumulation lines. Because UV-light was not used during the experiment, we are able to demonstrate that TCR in flies is not only limited to UV-induced damage, as previously seen (17), but can also repair other types of DNA damage. The C>A mutations can arise from mismatches with oxidatively damaged DNA (35). An important finding is that TCR does not cause mutational asymmetry in exons. We ruled out that this is caused by the MMR deficiency and found support for a pathway that protects exons over introns thus masking the signatures of TCR. A similar finding was made in human cells where less oxidative DNA damage accumulates in exons than in introns — possibly due to a favorable repair (33). If a similar process is occurring in flies, as our data suggest, we propose that the global repair pathway of nucleotide excision repair is favoring exons over introns. The global repair does not discriminate between the transcribed and non-transcribed strands and detects the same lesions as TCR thus explaining the lack of strand bias, the gene expression difference, and the signs of reduced exonic mutation rate on the non-transcribed strand for C>A mutations.

In summary, generating the largest de novo mutation data set from an MA study in *D. melanogaster*, we demonstrated that TCR operates against DNA damage in the germline independent of the MMR pathway. We uncovered differences in mutational processes of exons and introns and attribute this to an additional repair operating on exons. We anticipate the use of *spel1* mutations will become a widely used approach to study mutation patterns in a broad range of species.

## Materials and methods

### Generating the *spel1* deletion and mutation accumulation

The *spel1* null mutant was generated from an isogenized *Oregon-R* strain using the CRISPR-Cas9 genome engineering tool. The 2^nd^ and the 3^rd^ chromosomes were isogenized with balancers and the variation on the X chromosome was reduced by 5 generations of full-sib mating. Two gRNAs targeting the second and the last exon of *spel1* were cloned with the Gibson Assembly® Cloning Kit (New England Biolabs) into a BbsI (10,000 units/ml, NEB, R0539) digested pCDF4 (50) (Addgene plasmid # 49411; http://n2t.net/addgene:49411; RRID:Addgene_49411) expression vector. The ligation product was transformed into SURE2 cells and the construct was verified by Sanger sequencing.

A template for homology-directed repair was generated by Golden gate cloning. 1 kb homology arms were amplified from genomic DNA with primers LT41-LT44. Purified amplicons were mixed (30 ng each) with 50 ng pJET1.2-STOP-dsRed (51) (Addgene plasmid # 60944; http://n2t.net/addgene:60944; RRID:Addgene_60944), 50 ng pBS-GGAC-ATGC (51) (Addgene plasmid # 60949; http://n2t.net/addgene:60949; RRID:Addgene_60949), 1.5 μl 10x T4 ligation buffer, 1 μl BsmBI (10,000 units/ml, NEB, R0580), and water was added to 14 μl. After incubation for min at 55°C 1 μl T4 ligase (400,000 units/ml, NEB, M0202) was added. Ligation was performed by cycling the reaction between 5 min in 42°C and 5 min in 16°C overnight. Final digestion was performed for 30 min in 55°C followed by 10 min at 80°C to inactivate the enzyme. The ligation product was treated with Plasmid-safe nuclease (10,000 units/ml, Epicentre, E3101K) and transformed into SURE2 cells. Positive colonies were identified with colony PCR and recovered plasmids were verified by sequencing.

The germline transformation was achieved by microinjecting a mixture of the template (500 ng/μl), the gRNA expression vector (100 ng/μl), and pHsp70-Cas9 (52) (250 ng/μl) (Addgene plasmid # 60944; http://n2t.net/addgene:60944; RRID:Addgene_60944) into dechorionated fly embryos. F1 progeny were screened for the 3XP3::DsRed marker and a correct targeting of *spel1* was confirmed with PCR and sequencing. A PCR was performed to detect the double integration where the template plasmid integrates twice into the locus with the backbone. All the primers used in this study are listed in Supplementary Table 3.

We performed 10 generations of mutation accumulation with full-sib mating and sequenced individual females from 7 surviving lines.

### Library preparation and sequencing

Genomic DNA was extracted from a single female fly of each MA line using a standard high salt extraction method (36) with RNase A treatment. From each female, 70 ng genomic DNA was used to prepare paired-end libraries with the NEBNext Ultra II FS DNA Library Prep Kit (New England Biolabs, Ipswich, MA) using only 10% of the reagents recommended in the original protocol of the supplier. After double sided size selection targeting an insert size of 300 bp, libraries were amplified with dual-index primers using 5 PCR cycles. After purification with AMPureXP beads (Beckman Coulter, Brea, CA), the 7 libraries were quantified using the Qubit dsDNA HS Kit (Invitrogen, Carlsbad, CA), combined in equimolar amounts with additional 4 libraries from another experiment and sequenced on one lane of a HiSeq2500 using a 2×125bp protocol.

### QC and reads mapping

Libraries were first demultiplexed using ReadTools (37) (version 1.5.2; AssignReadGroupByBarcode --splitSample, --maximumMismatches 1, providing the corresponding barcodes). The raw reads were assessed for their quality using FastQC software (http://www.bioinformatics.babraham.ac.uk/projects/fastqc/). Low quality tails at 3’ end were trimmed using ReadTools (--mottQualityThreshold 20, --minReadLength 50, --disable5pTrim true) and BAM files were converted to compressed FASTQ files using ReadTools (ReadsToFastq -- interleavedInput true --barcodeInReadName true --outputFormat GZIP). As FastQC detected residual levels of adapter contamination, adapter cleaning was performed with the BBTools suit (38) using BBDuk (version 38.32; ktrim=r k=23 mink=11 hdist=1 tbo).

Processed paired-end reads were mapped to the *D. melanogaster* reference genome release 6.24 indexed with the bwa index command using BWA-MEM (39) (version 0.7.17; bwamem) on a Hadoop cluster using DistMap (40) (version 2.7.5).

PCR duplicates were removed using PICARD (http://broadinstitute.github.io/picard/) MarkDuplicates tool (version 2.21.3; REMOVE_DUPLICATES=true VALIDATION_STRINGENCY=SILENT). We kept mapped reads with each segment properly aligned and removed reads that mapped equally well to multiple positions or have a low mapping quality using SAMtools (41) (version 1.9; -b -q 20 -f 0×002 -F 0×004 -F 0×008). We clipped the processed overlapping paired-end reads using the BamUtil suit (42) (version 1.0.13; bam clipOverlap --in --out --stats).

### Variants inventory

The fasta reference was indexed using SAMtools faidx command. Processed BAM files were sorted and indexed with SAMtools for each chromosome arm (2L, 2R, 3L, 3R, X) separately using SAMtools view command. We then added a unique read group tag per sample using the PICARD AddOrReplaceReadGroups command. We increased the accuracy of variant calling by using two different tools; Freebayes (45, cloned from https://github.com/ekg/freebayes) (version v0.9.10-3-g47a713e) and GATK HaplotypeCaller (46) (version 4.0.12.0) and kept only variants that were identified with both tools. To use the parallel version of Freebayes, we split the reference into 1Mb regions with Freebayes fasta_generate_regions.py script (python version 2.7.17). For each chromosome arm, we used the freebayes-parallel executable (-C 1 –F 0.01 --min-base-quality 20, all other options set to default), providing the 1 Mb regions file and individual BAM files. Second, we followed (31) and used the GATK HaplotypeCaller with --heterozygosity 0.01 option with default settings.

We obtained two raw lists of variants per chromosome arm in a VCF format (43). Two different filtering procedures were applied for each variant caller.

For Freebayes, each raw list of variants was filtered as follows:

i. to remove variants based on depth at the variant position using BCFtools (44) (version 1.8; filter –i SAF>0 && SAR>0 && (SAF+SAR+SRF+SRR)>5),
ii. to suppress variants within 5-bp of an INDEL using BCFtools (filter -g 5),
iii. to keep variants with at most 2 alleles denoted by reference and alternate alleles using VCFtools (43) (version 0.1.15; --vcf --min-alleles 1 --max-alleles 2 --recode-INFO-all --recode),
iv. to simplify multi-nucleotide polymorphisms into SNPs using vt cloned from https://github.com/atks/vt (45) (version 0.57721; decompose_blocksub, normalize commands successively),
v. to filter for QUAL>40 using VCFtools (--vcf –minQ 40 --recode-INFO-all --recode).

For GATK, we used GATK VariantFiltration with the options --filter-expression “QD < 2.0” --filter-name “QD” --filter-expression “FS > 60.0” --filter-name “FS” --filter-expression “MQ < 40.0” --filter-name “MQ” --filter-expression “MQRankSum < -12.5” --filter-name “MQRankSum” --filter-expression “ReadPosRankSum < -8.0” --filter-name “ReadPosRankSum”.

We intersected the two filtered VCF files retaining only variants with the same position using BEDtools (46) (version 2.27.1; intersect –u –a -b -wa –header). We then extracted private SNPs using BEDtools (intersect –v –a -b –header) providing all bgzipped and tabix-indexed (47) (version 1.8; -p vcf) VCF per line. Finally, we subtracted the variants lists with the variants called from 10 individual *spel1* null flies, which did not go through MA, as a quality control for residual ancestral alternative alleles after having applied a similar pipeline; we masked the X region 6240639:6686943 from line 5 using BEDtools (intersect –v –a -b –header) where some residual variants were observed. We obtained a final set of 7,345 SNPs and 5,672 INDELs.

For our analyses we relied on the genome annotation from flybase Dmel-all-filtered-r6.30.gff (downloaded from: ftp://ftp.flybase.net/genomes/Drosophila_melanogaster/dmel_r6.30_FB2019_05/gff/ in May 2019).

### Statistical analyses

All statistical analyses were done with R (48) (version 3.5.0).

#### Fraction of non-synonymous mutations compared to neutral expectation

The SnpEff software (49) (version 4.3) was used to distinguish synonymous and nonsynonymous mutations in the longest transcript of each gene. We performed a Fisher’s exact test to compare the observed and expected number of synonymous and non-synonymous mutations. Following (31), we used odds of 1:3 for synonymous and non-synonymous mutations as a neutral expectation.

#### Skew of intronic base composition

Gene expression data from ovaries and testes tissues were obtained from FlyAtlas2 (34), representing 16,781 genes. FPKM gene expression values were grouped into 40 bins, separately for ovaries and testes with the mltools::bin_data (50) (version 0.3.5; binType=“quantile”) R function. Since alternative splicing may generate ambiguous signals, 7 bases from the 5’ end and 35 bases from the 3’ end were removed from introns to exclude genomic regions containing splicing sequences as recommended in (15). AT (CG) skews were then calculated as the number of T (C) on the transcribed strand over the total number of A and T (C and G) bases. For each tissue and type of skew, we fitted a Generalized Linear Model (51) using the glm(cbind(#transcribed, #total-#transcribed), family=“binomial”) R function, and reported the Wald test p-values corresponding to the binned gene expression covariate.

#### Mutational strand bias

We restricted our analysis to unambiguous exons and introns and excluded annotations overlapping with other genes located on a different strand using the BEDtools intersect -s command (a GTF with the final annotation can be found in the Dryad repository). We used the Bioconductor MutationalPatterns package (52) (version 1.12.0) to count the different mutation types on the transcribed and non-transcribed strands. Our first approach was to estimate the expected mutation rate from the base composition on the transcribed and non-transcribed strand of genes with at least one mutation. Since without strand bias a ratio of 1 is expected, we calculated its significance and 95% confidence intervals using the poisson.test R function.

In the second approach, we accounted for the impact of gene expression intensity on base composition. For each of the 40 expression bins, we randomly sampled the same number of genes as observed being mutated in our SNPs set and calculated the expected strand bias from the sample.

We repeated the approaches for the expected intronic and exonic biases, using exclusively either intronic or exonic sequences. The p-values were corrected for multiple testing using the Benjamini-Hochberg procedure.

In order to take the 5’ and 3’ flanking bases of the A[T>C]N mutations into account in the null hypothesis, we adapted a permutation procedure from (32) to test for strand bias in exons and introns (Supplementary Figure 3). Briefly, we obtained the frequency of mutations for each of the 4 A[T>C]N contexts (triplets) genome-wide and rescaled the frequencies to sum up to 1. In parallel, we used the GATK tool CallableLoci (53) to obtain the callable sites per line and the BEDtools suit (maskfasta and getfasta commands) to mask the reference for non-callable sites. For both strands, we then retained as a sampling pool the number of callable triplets in the mutated genes for exons, introns, summed over each line, and multiplied it with the rescaled frequency to weight the sampling according to the genome-wide prevalence of triplets. Finally, we redistributed the observed number of mutations on the transcribed and non-transcribed strand separately 10,000 times to get the expected number of mutations on the transcribed strand in introns and exons. The p-values were calculated as the number of times the sampled value was higher than the observed one divided by 10,000.

#### Gene expression analysis for C>A and A[T>C]N mutations

Gene expression differences between genes containing C>A and A[T>C]N mutations on different strands were tested with either one-sided (intron) or two-sided (exon) Wilcoxon rank-sum test on the FPKM scale. We used a one-sided test for intronic sequences because the strand bias predicts the direction of gene expression difference. The p-values were corrected for multiple testing using the Benjamini-Hochberg procedure.

#### Mutagenic effect of transcription

To test if transcription is mutagenic, we performed a randomization test similarly as (54) (Supplementary Figure 4). We randomly picked 221 genes, corresponding to the number of C>A intronic mutations overlapping with the FlyAtlas2 data (over 236) on the non-transcribed strand, and computed the mean expression in ovaries and testes separately. The sampling was weighted by the length of the introns. This was done 10,000 times. For each tissue, a p-value was calculated as the number of times the randomly sampled mean values exceeded the observed mean divided by 10,000.

#### Decreased exonic mutation rate for C>A and A[T>C]N mutations

We used a similar permutation procedure as described above in the mutational strand bias subsection to test for reduced exonic mutation rates (Supplementary Figures 5, 6). We modified the sampling pool of callable triplets to include the genome-wide exons and introns with strands separated.

### Code and data availability

The code (R and bash scripts) will be accessible in the following github repository: ***, available upon publication.

The final set of SNPs and INDELs as well as the updated annotation and intermediate files can be found from the following dryad repository: ***, available upon publication.

Raw reads will be available in the following SRA project: ***, available upon publication.

## Authors contribution

L. T. performed experiments, V. N. performed sequencing, L. T., C. B. analyzed the data, L. T., C. B., V. N., C. S. wrote the paper, K.S. supervised the project and provided feedback, L. T., C. S. designed the study.

## Acknowledgments

Sequencing was performed at the VBCF NGS Unit (https://www.viennabiocenter.org/facilities/next-generation-sequencing/). This work was supported by the Austrian Science Fund (FWF, grants W1225, P27630, P29133). We thank Lukas Endler for technical comments.

---

**Supplementary Figure 1.**
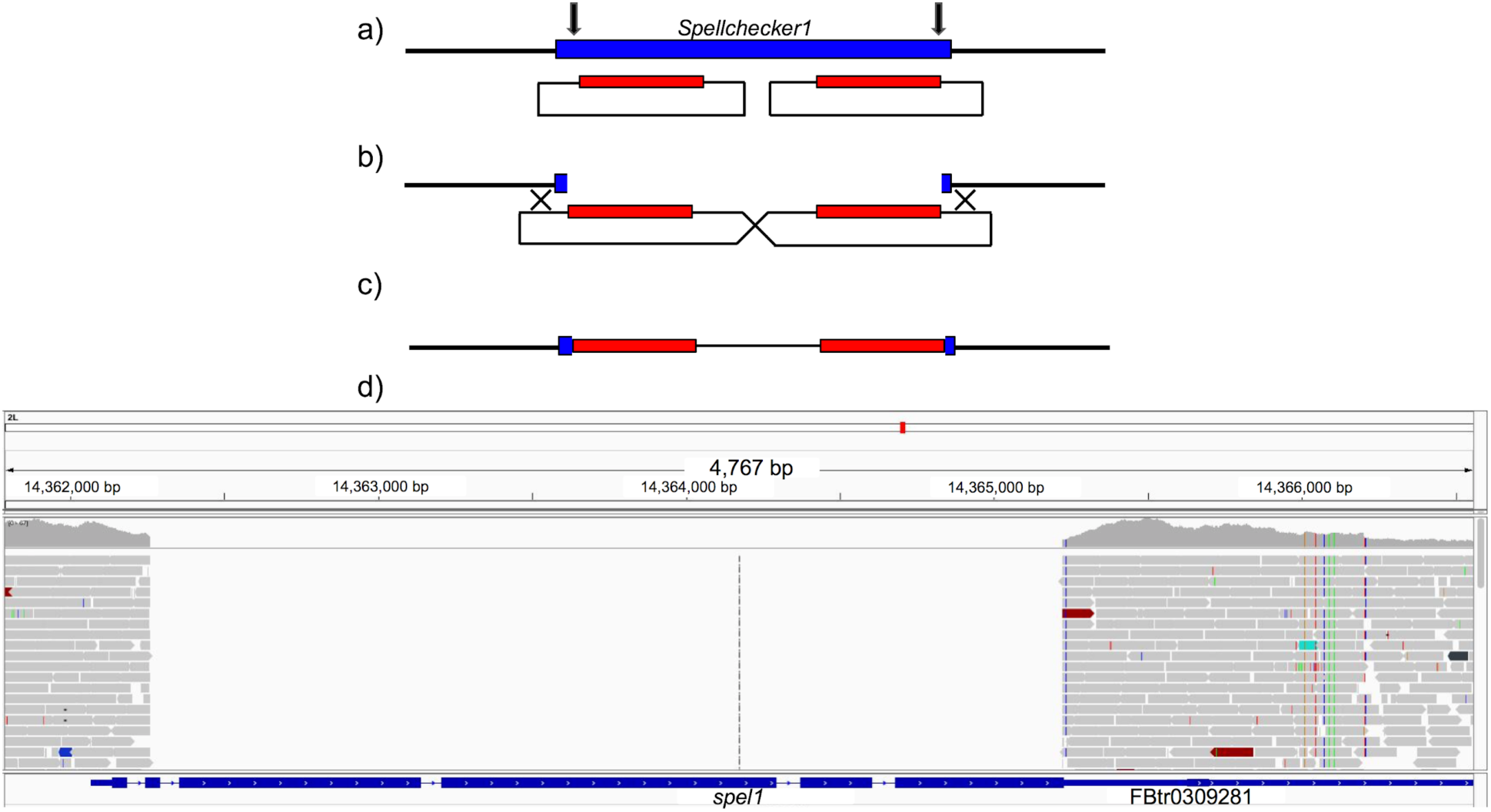
A schematic overview showing the double insertion of the template plasmid and the short Illumina-read based confirmation of the *spel1* deletion. a) Two gRNAs targeting the gene are indicated by black arrows b) the resulting double-stranded break is repaired by two template plasmids which recombine with each other c) the resulting allele contains the backbone of the plasmid flanked by dsRed cassettes d) a screenshot of the short read coverage at the *spel1* locus visualized by IGV (55).

**Supplementary Figure 2.**
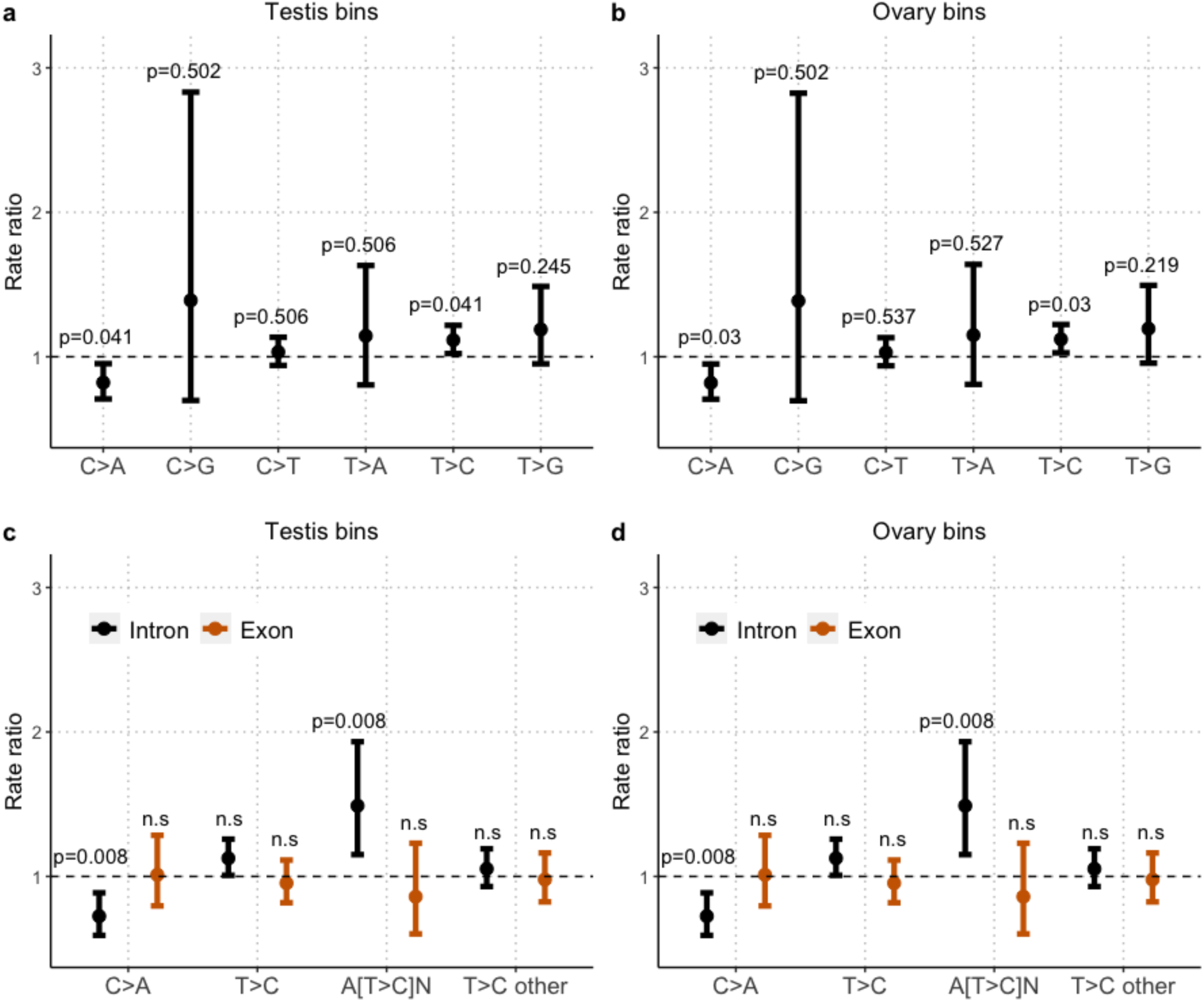
Mutational bias within genes tested using the gene expression sampling scheme. Estimated rate ratios (RR) on the transcribed strand by the non-transcribed strands (dot) in genes and partitioned in d) introns (black) and exons (orange) based on testis (a, c) and b) ovary (b, d) expression bins. Adjusted p-values using the Benjamini-Hochberg procedure are reported and 95% Poisson confidence intervals are represented by segments. The intercept line of 1 indicates the absence of differences in rates of mutations on the transcribed and non-transcribed strands. The significance threshold is set to 5%.

**Supplementary Figure 3.**
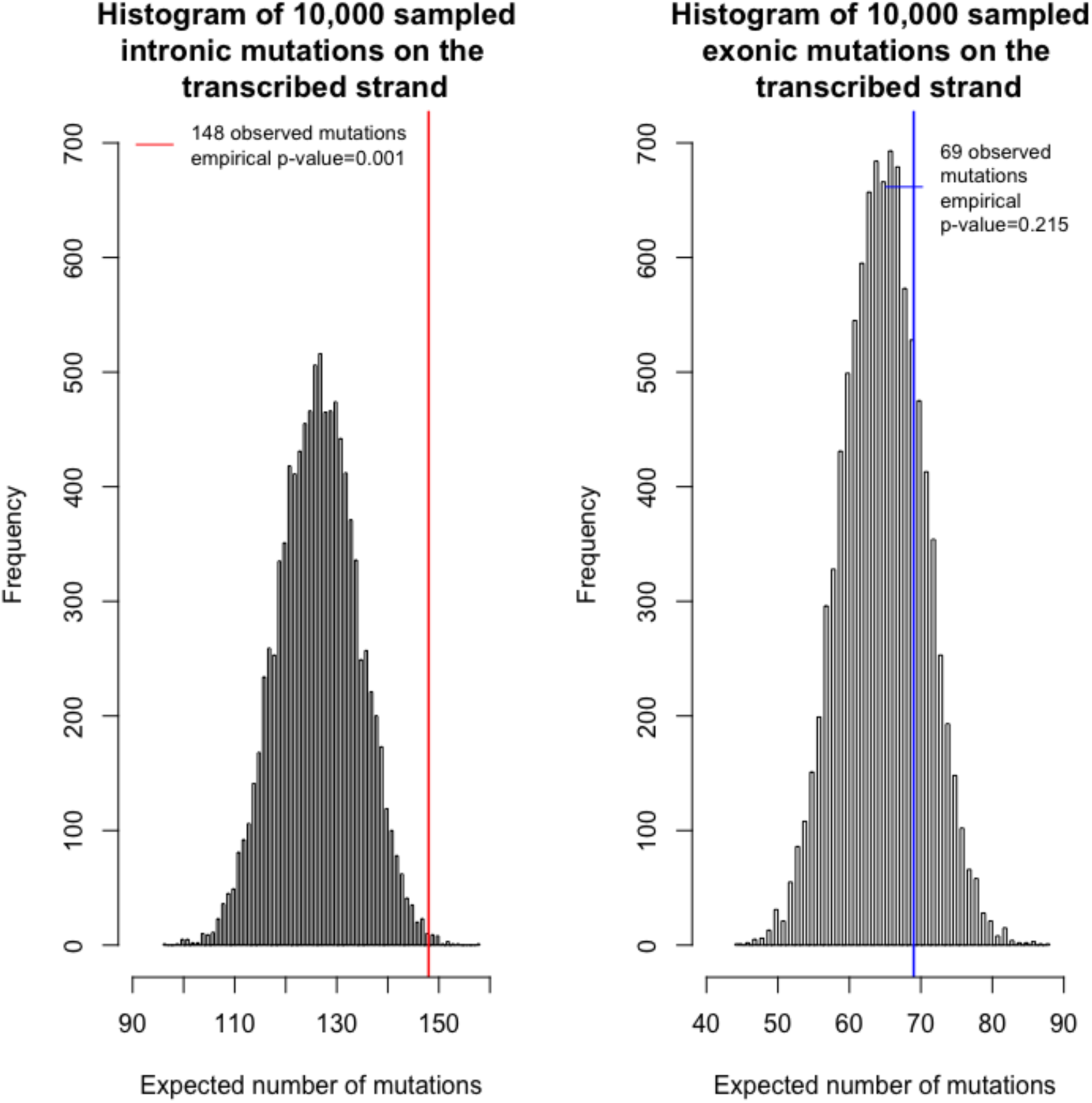
Permutation test to test strand bias for A[T>C]N mutations while taking the composition of the 5’ and 3’ flanking bases into account. The histograms show the expected number of A[T>C]N on the transcribed strand in introns (left) and exons (right). The red line shows the observed mutations on the transcribed strand in introns and the blue line exons.

**Supplementary Figure 4.**
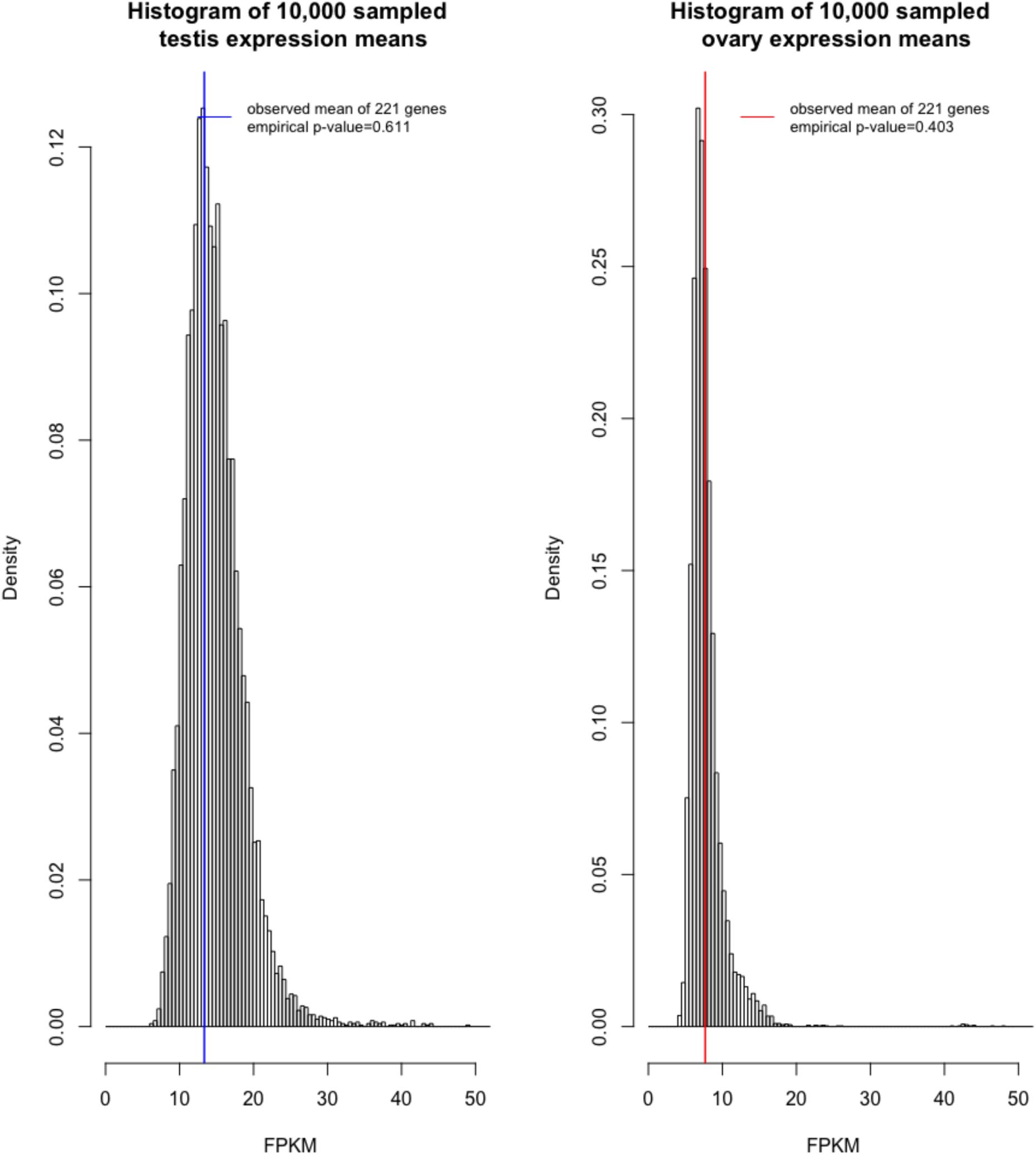
Randomization test to rule out the mutagenic effect of transcription. Histograms show the frequency distribution of the mean expression level of 221 randomly sampled genes in testes (left) and ovaries (right) from 10,000 trials. Vertical lines show the observed mean expression of genes with C>A mutations located on the non-transcribed strand. The binning is the same between both histograms.

**Supplementary Figure 5.**
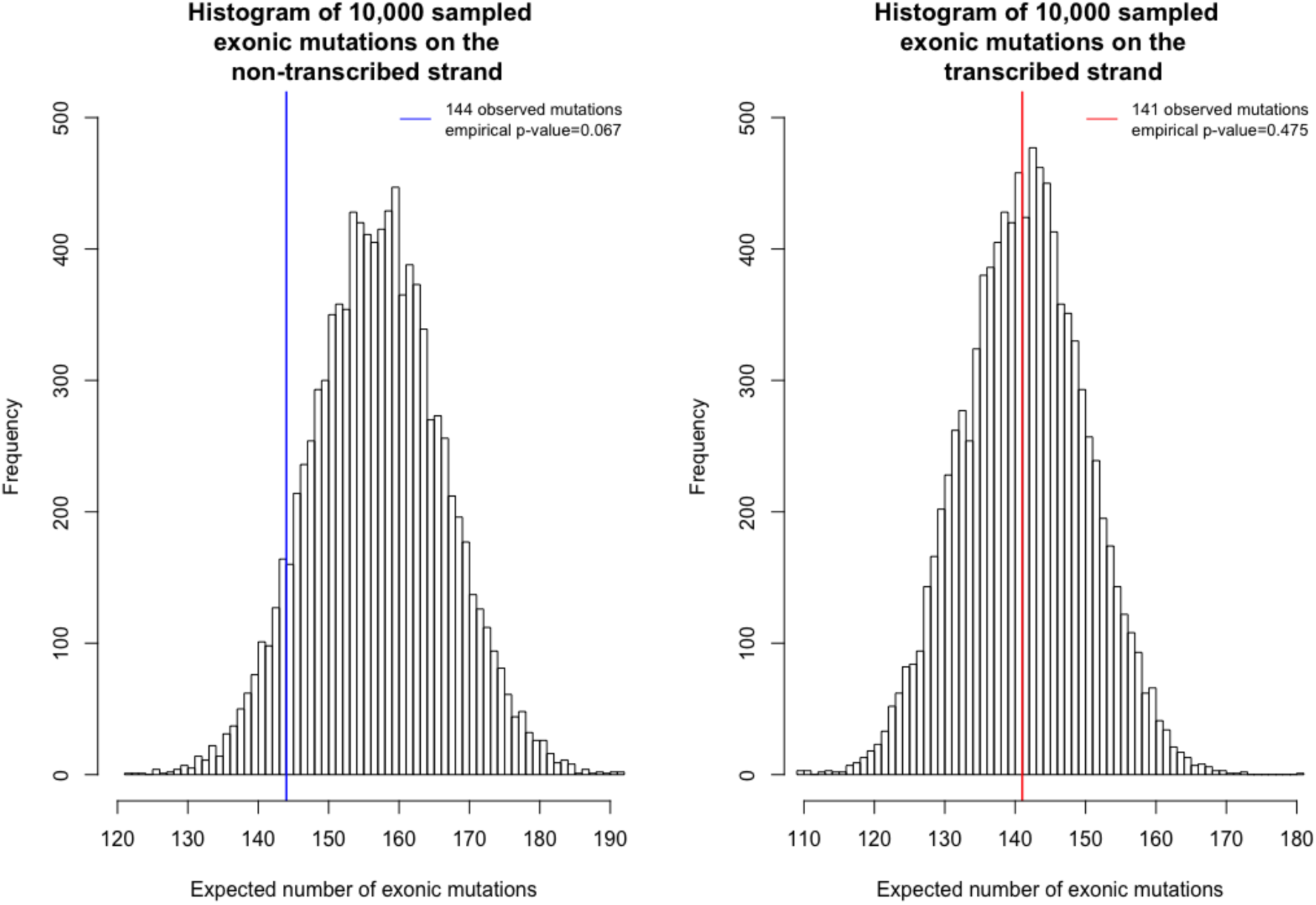
Permutation test to test for reduced exonic C>A mutation rate. Histograms show the sampled number of exonic mutations when the expectation was calculated based on the non-transcribed strand (left) and transcribed strand (right). Vertical lines show the observed number for C>A mutations when the cytosine was located on the non-transcribed strand (blue) and on the transcribed strand (red).

**Supplementary Figure 6.**
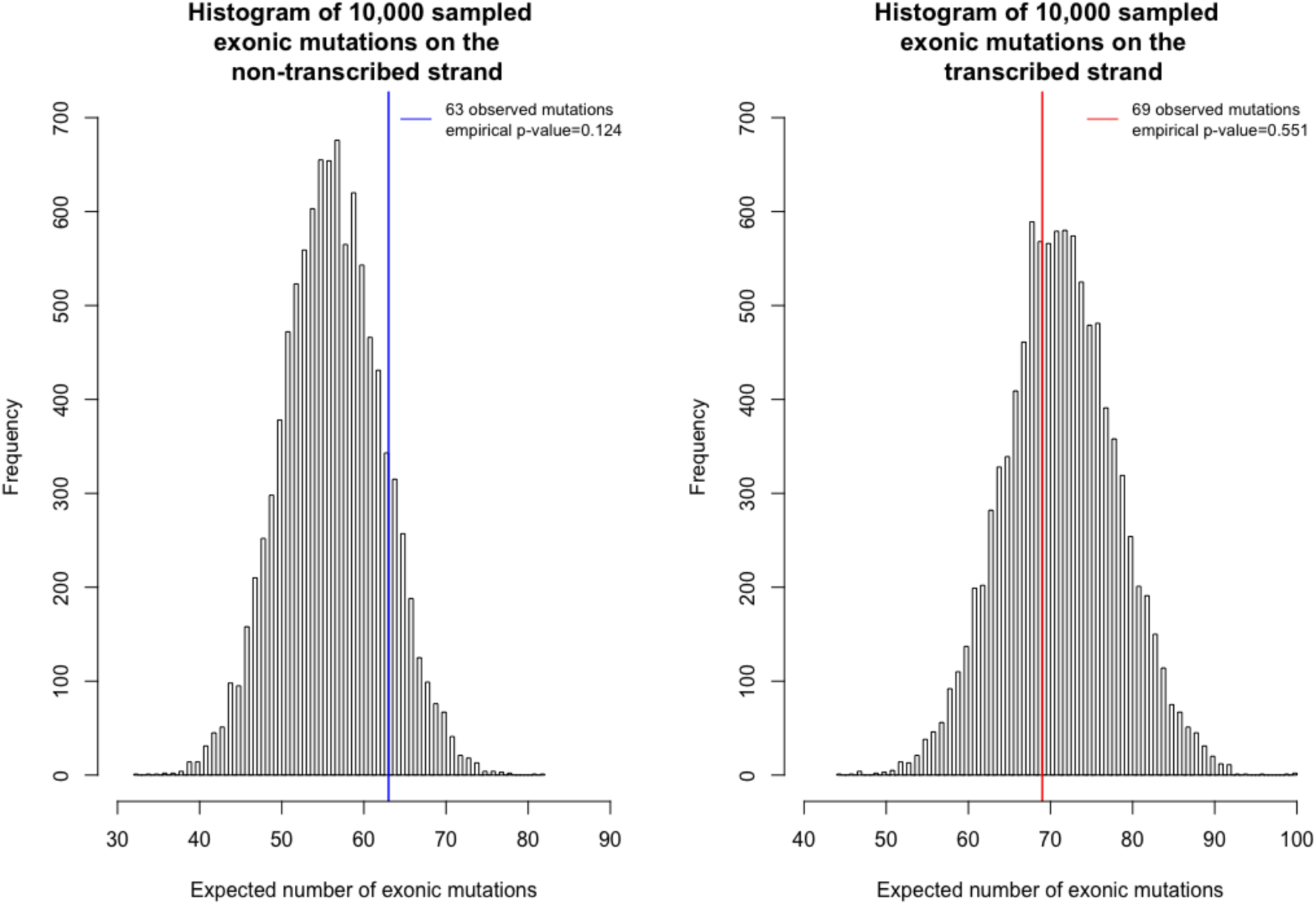
Permutation test to test for reduced exonic A[T>C]N mutation rate. Histograms show the sampled number of exonic mutations when the expectation was calculated based on the non-transcribed strand (left) and transcribed strand (right). Vertical lines show the observed number for A[T>C]N mutations when the thymine was located on the non-transcribed strand (blue) and on the transcribed strand (red).

**Supplementary Table 1.** Observed SNV counts in genes. Raw p-values were obtained using a Poisson test and corrected with a Benjamini-Hochberg procedure.

| Type | Transcribed | Non-transcribed | Total | Rate ratio (95% CI) | Adjusted p-value |
| --- | --- | --- | --- | --- | --- |
| C>A | 329 | 405 | 734 | 0.815 (0.702-0.945) | 0.0376 |
| C>G | 22 | 16 | 38 | 1.379 (0.692-2.809) | 0.5041 |
| C>T | 896 | 877 | 1,773 | 1.024 (0.933-1.126) | 0.6180 |
| T>A | 72 | 63 | 135 | 1.145 (0.805-1.632) | 0.5279 |
| T>C | 1087 | 976 | 2,063 | 1.116 (1.023-1.218) | 0.0385 |
| T>G | 178 | 150 | 328 | 1.189 (0.951-1.488) | 0.2444 |

**Supplementary Table 2.** Observed SNV counts in introns and exons. Raw p-values were obtained using a Poisson test and corrected using the Benjamini-Hochberg procedure.

| Type | Transcribed | Non-transcribed | Total | Rate ratio (95% CI) | Adjusted p-value | Annotation |
| --- | --- | --- | --- | --- | --- | --- |
| C>A | 171 | 236 | 407 | 0.732 (0.597-0.895) | 0.00982 | Intron |
| T>C | 667 | 620 | 1287 | 1.112 (0.996-1.243) | 0.15454 | Intron |
| A[T>C]N | 148 | 104 | 252 | 1.472 (1.138-1.910) | 0.00982 | Intron |
| T>C Other | 519 | 516 | 1,035 | 1.040 (0.919-1.178) | 0.86566 | Intron |
| C>A | 141 | 144 | 285 | 0.994 (0.783-1.264) | 1 | Exon |
| T>C | 373 | 307 | 680 | 0.993 (0.851-1.158) | 1 | Exon |
| A[T>C]N | 69 | 63 | 132 | 0.894 (0.627-1.280) | 0.86566 | Exon |
| T>C Other | 304 | 244 | 548 | 1.018 (0.857-1.210) | 1 | Exon |

**Supplementary Table 3.**
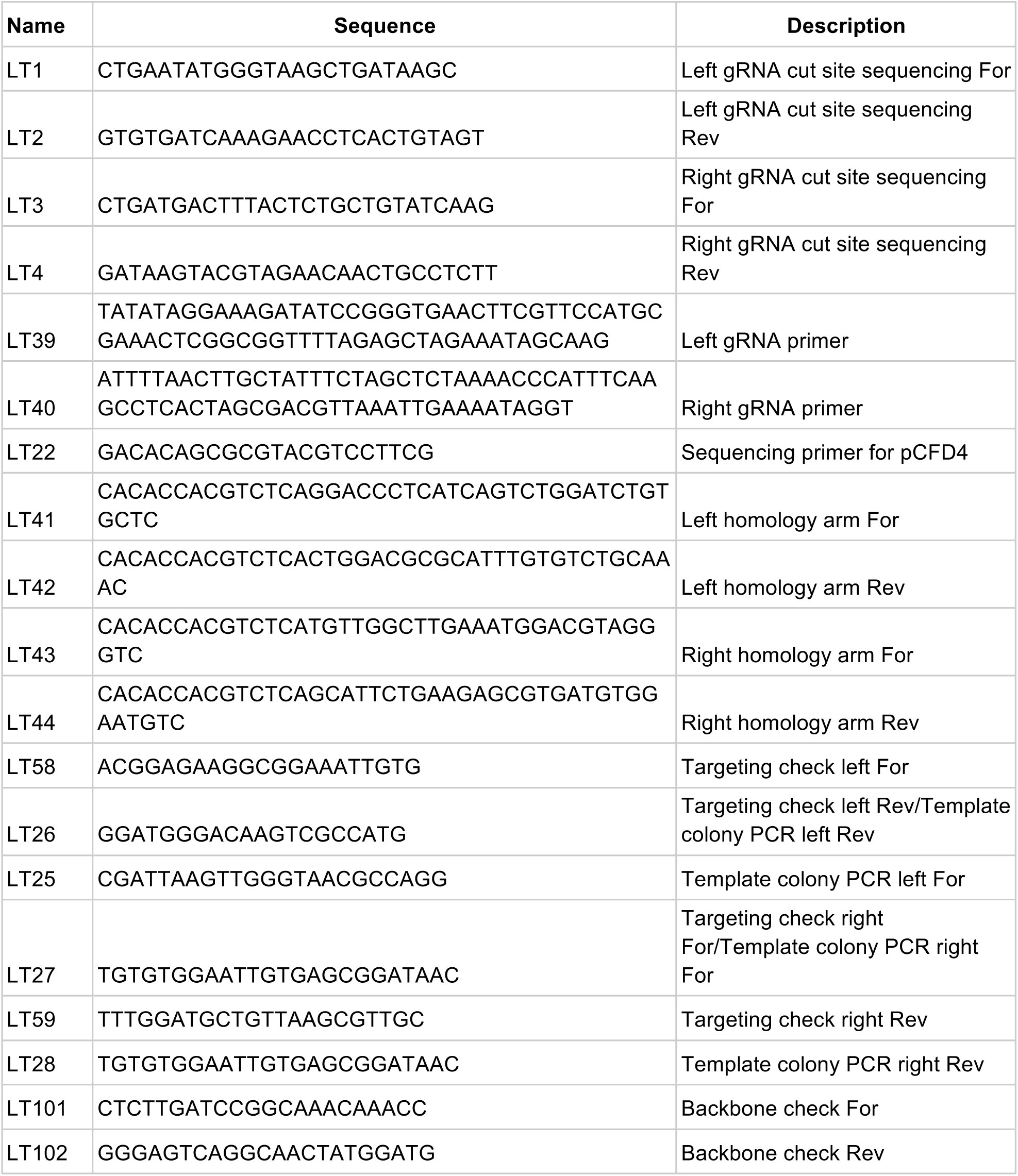
Primers used to generate the *sple1* null mutant.

## References

1. T. Lindahl, Instability and decay of the primary structure of DNA. Nature 362, 709–715 (1993).

2. J. Nakamura, et al., Highly sensitive apurinic/apyrimidinic site assay can detect spontaneous and chemically induced depurination under physiological conditions. Cancer Res. 58, 222–225 (1998).

3. W. Wang, C. Walmacq, J. Chong, M. Kashlev, D. Wang, Structural basis of transcriptional stalling and bypass of abasic DNA lesion by RNA polymerase II. Proc. Natl. Acad. Sci. U. S. A. 115, E2538–E2545 (2018).

4. J. A. Marteijn, H. Lans, W. Vermeulen, J. H. J. Hoeijmakers, Understanding nucleotide excision repair and its roles in cancer and ageing. Nat. Rev. Mol. Cell Biol. 15, 465–481 (2014).

5. P. C. Hanawalt, G. Spivak, Transcription-coupled DNA repair: two decades of progress and surprises. Nat. Rev. Mol. Cell Biol. 9, 958–970 (2008).

6. P. Green, et al., Transcription-associated mutational asymmetry in mammalian evolution. Nat. Genet. 33, 514–517 (2003).

7. L. E. Kerry, et al., Selective inhibition of RNA polymerase I transcription as a potential approach to treat African trypanosomiasis. PLoS Negl. Trop. Dis. 11, e0005432 (2017).

8. A. Zheleva, D. Michelot, Z. D. Zhelev, Sensitivity of alpha-amanitin to oxidation by a lactoperoxidase-hydrogen peroxide system. Toxicon 38, 1055–1063 (2000).

9. J. A. Eisen, P. C. Hanawalt, A phylogenomic study of DNA repair genes, proteins, and processes. Mutat. Res. 435, 171–213 (1999).

10. A. R. Lehmann, The xeroderma pigmentosum group D (XPD) gene: one gene, two functions, three diseases. Genes Dev. 15, 15–23 (2001).

11. J. J. Sekelsky, M. H. Brodsky, K. C. Burtis, DNA repair in Drosophila: insights from the Drosophila genome sequence. J. Cell Biol. 150, F31–6 (2000).

12. J. Sekelsky, DNA Repair in Drosophila: Mutagens, Models, and Missing Genes. Genetics 205, 471–490 (2017).

13. J. G. de Cock, et al., Repair of UV-induced (6-4)photoproducts measured in individual genes in the Drosophila embryonic Kc cell line. Nucleic Acids Res. 20, 4789–4793 (1992).

14. P. J. van der Helm, E. C. Klink, P. H. Lohman, J. C. Eeken, The repair of UV-induced cyclobutane pyrimidine dimers in the individual genes Gart, Notch and white from isolated brain tissue of Drosophila melanogaster. Mutat. Res. 383, 113–124 (1997).

15. M. Touchon, A. Arneodo, Y. d’Aubenton-Carafa, C. Thermes, Transcription-coupled and splicing-coupled strand asymmetries in eukaryotic genomes. Nucleic Acids Res. 32, 4969–4978 (2004).

16. J. Bergman, A. J. Betancourt, C. Vogl, Transcription-Associated Compositional Skews in Drosophila Genes. Genome Biol. Evol. 10, 269–275 (2018).

17. N. Deger, Y. Yang, L. A. Lindsey-Boltz, A. Sancar, C. P. Selby, Drosophila, which lacks canonical transcription-coupled repair proteins, performs transcription-coupled repair. J. Biol. Chem. 294, 18092–18098 (2019).

18. I. Mellon, G. N. Champe, Products of DNA mismatch repair genes mutS and mutL are required for transcription-coupled nucleotide-excision repair of the lactose operon in Escherichia coli. Proc. Natl. Acad. Sci. U. S. A. 93, 1292–1297 (1996).

19. P. Bertrand, D. X. Tishkoff, N. Filosi, R. Dasgupta, R. D. Kolodner, Physical interaction between components of DNA mismatch repair and nucleotide excision repair. Proc. Natl. Acad. Sci. U. S. A. 95, 14278–14283 (1998).

20. K. S. Sweder, et al., Mismatch repair mutants in yeast are not defective in transcription-coupled DNA repair of UV-induced DNA damage. Genetics 143, 1127–1135 (1996).

21. J. Zhao, A. Jain, R. R. Iyer, P. L. Modrich, K. M. Vasquez, Mismatch repair and nucleotide excision repair proteins cooperate in the recognition of DNA interstrand crosslinks. Nucleic Acids Res. 37, 4420–4429 (2009).

22. A. Bellacosa, Functional interactions and signaling properties of mammalian DNA mismatch repair proteins. Cell Death Differ. 8, 1076–1092 (2001).

23. K. Kobayashi, P. Karran, S. Oda, K. Yanaga, The involvement of mismatch repair in transcription coupled nucleotide excision repair. Hum. Cell 18, 103–115 (2005).

24. G. Rossetti, et al., The structural impact of DNA mismatches. Nucleic Acids Res. 43, 4309–4321 (2015).

25. D. Mu, et al., Recognition and repair of compound DNA lesions (base damage and mismatch) by human mismatch repair and excision repair systems. Mol. Cell. Biol. 17, 760–769 (1997).

26. T. L. DeWeese, et al., Mouse embryonic stem cells carrying one or two defective Msh2 alleles respond abnormally to oxidative stress inflicted by low-level radiation. Proc. Natl. Acad. Sci. U. S. A. 95, 11915–11920 (1998).

27. C. Colussi, et al., The mammalian mismatch repair pathway removes DNA 8-oxodGMP incorporated from the oxidized dNTP pool. Curr. Biol. 12, 912–918 (2002).

28. V. Grazielle-Silva, et al., Distinct Phenotypes Caused by Mutation of MSH2 in Trypanosome Insect and Mammalian Life Cycle Forms Are Associated with Parasite Adaptation to Oxidative Stress. PLoS Negl. Trop. Dis. 9, e0003870 (2015).

29. F. Le Page, A. Klungland, D. E. Barnes, A. Sarasin, S. Boiteux, Transcription coupled repair of 8-oxoguanine in murine cells: the ogg1 protein is required for repair in nontranscribed sequences but not in transcribed sequences. Proc. Natl. Acad. Sci. U. S. A. 97, 8397–8402 (2000).

30. Y. Ding, A. Berrocal, T. Morita, K. D. Longden, D. L. Stern, Natural courtship song variation caused by an intronic retroelement in an ion channel gene. Nature 536, 329–332 (2016).

31. Z. J. Assaf, S. Tilk, J. Park, M. L. Siegal, D. A. Petrov, Deep sequencing of natural and experimental populations of Drosophila melanogaster reveals biases in the spectrum of new mutations. Genome Res. 27, 1988–2000 (2017).

32. J. Frigola, et al., Reduced mutation rate in exons due to differential mismatch repair. Nat. Genet. 49, 1684–1692 (2017).

33. A. R. Poetsch, S. J. Boulton, N. M. Luscombe, Genomic landscape of oxidative DNA damage and repair reveals regioselective protection from mutagenesis. Genome Biol. 19, 215 (2018).

34. D. P. Leader, S. A. Krause, A. Pandit, S. A. Davies, J. A. T. Dow, FlyAtlas 2: a new version of the Drosophila melanogaster expression atlas with RNA-Seq, miRNA-Seq and sex-specific data. Nucleic Acids Res. 46, D809–D815 (2018).

35. S. A. Lujan, et al., Heterogeneous polymerase fidelity and mismatch repair bias genome variation and composition. Genome Res. 24, 1751–1764 (2014).

36. S. A. Miller, D. D. Dykes, H. F. Polesky, A simple salting out procedure for extracting DNA from human nucleated cells. Nucleic Acids Res. 16, 1215 (1988).

37. D. Gómez-Sánchez, C. Schlötterer, ReadTools: A universal toolkit for handling sequence data from different sequencing platforms. Mol. Ecol. Resour. 18, 676–680 (2018).

38. B. Bushnell, BBMap. sourceforge.

39. H. Li, R. Durbin, Fast and accurate short read alignment with Burrows-Wheeler transform. Bioinformatics 25, 1754–1760 (2009).

40. R. V. Pandey, C. Schlötterer, DistMap: a toolkit for distributed short read mapping on a Hadoop cluster. PLoS One 8, e72614 (2013).

41. H. Li, et al., The Sequence Alignment/Map format and SAMtools. Bioinformatics 25, 2078–2079 (2009).

42. M. R. Breese, Y. Liu, NGSUtils: a software suite for analyzing and manipulating next-generation sequencing datasets. Bioinformatics 29, 494–496 (2013).

43. P. Danecek, et al., The variant call format and VCFtools. Bioinformatics 27, 2156–2158 (2011).

44. H. Li, A statistical framework for SNP calling, mutation discovery, association mapping and population genetical parameter estimation from sequencing data. Bioinformatics 27, 2987–2993 (2011).

45. A. Tan, G. R. Abecasis, H. M. Kang, Unified representation of genetic variants. Bioinformatics 31, 2202–2204 (2015).

46. A. R. Quinlan, I. M. Hall, BEDTools: a flexible suite of utilities for comparing genomic features. Bioinformatics 26, 841–842 (2010).

47. H. Li, Tabix: fast retrieval of sequence features from generic TAB-delimited files. Bioinformatics 27, 718–719 (2011).

48. R Core Team, R: A Language and Environment for Statistical Computing (2018).

49. P. Cingolani, et al., A program for annotating and predicting the effects of single nucleotide polymorphisms, SnpEff: SNPs in the genome of Drosophila melanogaster strain w1118; iso-2; iso-3. Fly 6, 80–92 (2012).

50. B. Gorman, mltools: Machine Learning Tools.

51. P. McCullagh, Generalized linear models (Routledge, 2019).

52. F. Blokzijl, R. Janssen, R. van Boxtel, E. Cuppen, MutationalPatterns: comprehensive genome-wide analysis of mutational processes. Genome Med. 10, 33 (2018).

53. A. McKenna, et al., The Genome Analysis Toolkit: a MapReduce framework for analyzing next-generation DNA sequencing data. Genome Res. 20, 1297–1303 (2010).

54. C. Park, W. Qian, J. Zhang, Genomic evidence for elevated mutation rates in highly expressed genes. EMBO Rep. 13, 1123–1129 (2012).

55. J. T. Robinson, et al., Integrative genomics viewer. Nat. Biotechnol. 29, 24–26 (2011).

